# Ecological aspects of sandfly fauna (Diptera: Psychodidae, Phlebotominae) of Sumidouro District, State of Rio de Janeiro, Brazil

**DOI:** 10.1101/2020.07.22.212589

**Authors:** João Ricardo Carreira Alves, Cleber Nascimento do Carmo, Rodrigo Caldas Menezes, Mauricio Luís Vilela, Jacenir Reis dos Santos-Mallet

## Abstract

Aiming to compare and update the sand fly fauna of Portão de Pedra site, Sumidouro District, Rio de Janeiro State, Brazil, and considering the environmental changes occurred, the biology and ecology of the local sandfly species were examined five years later as a complementary study carried. Captures were made in the cave, surroundings of cave and forest of the region, from 6 p.m. to 6 a.m. 2 3 2 3 sandflies of eigth species of the *Lutzomyia* were captured: *L. gasparviannai, L. edwardsi, L. tupynambai, L. hirsuta, L. whitmani, L. migonei, L. intermedia, Lutzomyia*. sp and one species of the *Brumptomyia* Kind: *B. brumpti*. In 2009 and 2010 were collected 1756 samples from ten species of the former genus and two of the second. *L. gasparviannai* was predominant, in the three collection sites, in both periods. Five species implicated as vector of *Leishmania*: *L. intermedia, L. whitmani, L. migonei, L. hirsuta* and *L. davisi* have been collected in the area. Poisson regression and ANOVA were used to perform statistical analysis of species most relevant. The record of *L. intermedia* and a case of American tegumentary leishmaniasis are relevant to public health of municipality and of state of Rio de Janeiro.

## Introduction

About 544 species of sandflies are found in the Americas, of which 527 presently exist and 17 are fossils (Galati, 2018).

According to Young & Duncan (1994), Phlebotomine sandflies (Diptera, Psychodidae, Phlebotominae) are classified into three genera in the New World: *Warileya, Brumptomyia*, and *Lutzomyia*. The sandflies of the genus *Lutzomyia* França, 1924 (in the New World) are invertebrate hosts of species belonging to the genus *Leishmania* Ross, 1903, (Kinetoplastida, Trypanosomatidae), which cause leishmaniasis in humans and other mammals. These protozoans are transmitted through the bite of infected female sand flies (Rangel & Lainson, 2003).

In Brazil, 279 species of sandflies have been described so far, including 22 species that are either proven or suspected to transmit leishmaniasis agents (Aguiar & Vieira, 2018).

According to Aguiar & Vieira (2018), 125 species were recorded in the southeast region and 65 in the state of Rio de Janeiro (Carvalho et al. 2014), including 6 species that are considered to be *Leishmania* vectors for humans and mammals (Aguiar & Vieira, 2018).

After cases of American Tegumentary Leishmaniasis (ATL) were recorded in mountainous regions affected by human activity in the state of Rio de Janeiro, studies on sand fly fauna and their ecological aspects were initiated in these locations (Alves 2007 and 2008; Carreira-Alves 2008; De Souza et al., 1995; Peres-Dias et al., 2016; Souza et al., 2002; and Souza et al., 2003).

In January 2011, torrential rainfall in these mountainous regions, land use and occupation with cuts and embankments, previous rainfall, and river and rain erosion caused geological instability in the city of Sumidouro (SDEEIS, 2011).

Between 2007 and 2017, 31 cases of cutaneous leishmaniasis were recorded in Sumidouro and its neighboring municipalities (SINAN, 2019). Noteworthy is the discovery of a case of leishmaniasis that occurred in 2007, which had not been recorded by the local health authorities (Alves, personal communication) and the collection of *L. intermedia*, a vector of *Leishmania braziliensis*, was recorded for first time in 2015 and 2016.

This study, highlights the environmental changes that occurred in a neighborhood called São Caetano after five years, particularly in terms of biodiversity, population density, species dominance, and sand fly species that are *Leishmania* vectors.

Therefore, the results of analyses of the materials collected in 2009 and 2010 were compared with those in 2015 and 2016 in the same location in the city of Sumidouro.

## Material and methods

### Ethics statement

Collections were carried out in forest area, with a cave inside, located on a private property. A consent statement was established to perform the captures in this forest area and in the cave.

### Area of study

The city of Sumidouro is located at the latitude 22°02’59” south and longitude 42°40’29” west, in the mountain region of the State of Rio de Janeiro, with an altitude of 355 meters, bordering the cities of Nova Friburgo, Teresópolis, Carmo, São José do Vale do Rio Preto, Sapucaia and Duas Barras. It has a total area of 397.6 Km^2^, corresponding to 5.7% of the mountain region. It is located 174 km from Rio de Janeiro, capital of the state (PMS 2019). (Fig 1.)

**Fig 1.**
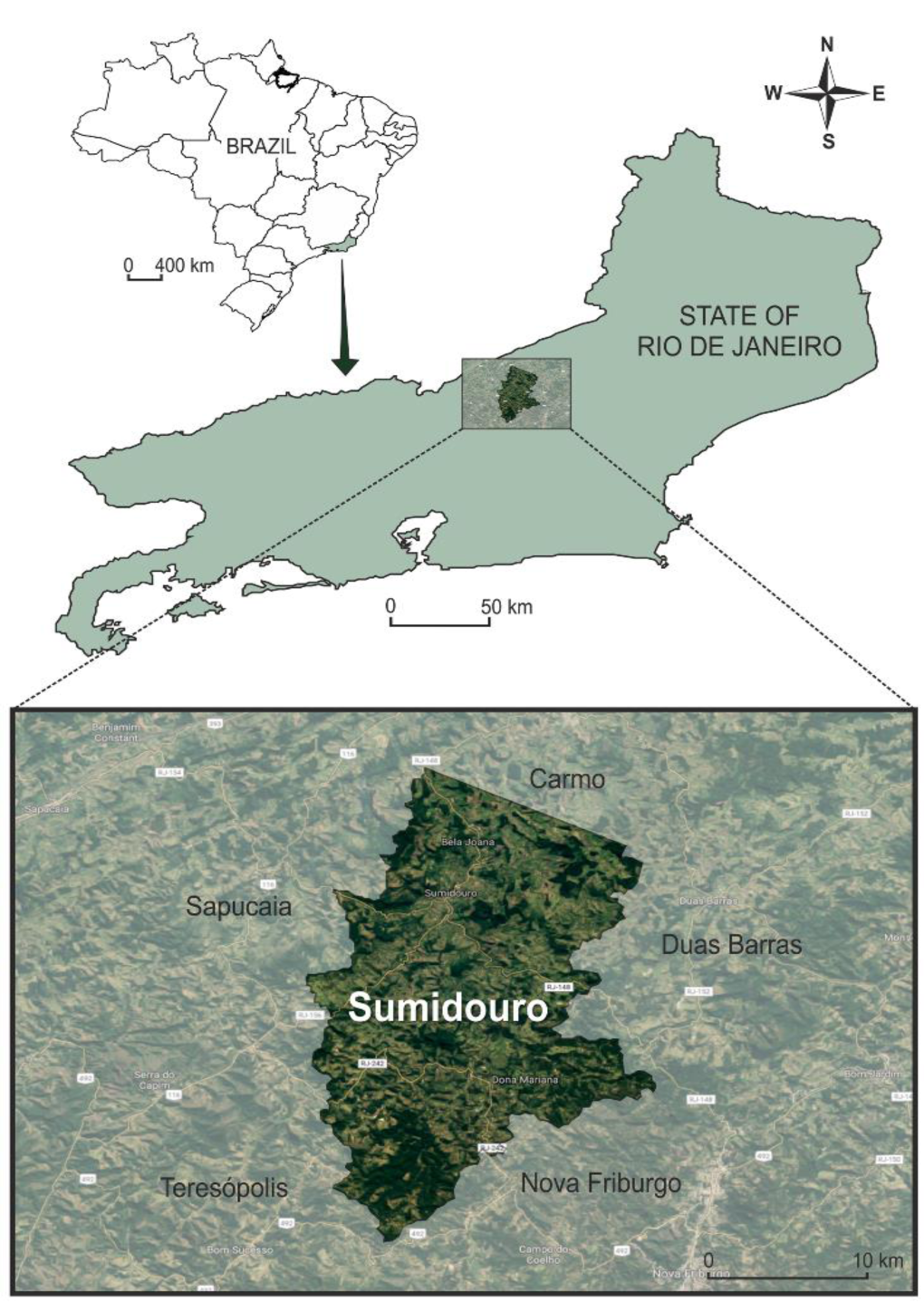
Geographic location of the city of Sumidouro, latitude 22°02’59” south and longitude 42°40’29” west, in the mountain region of the state of Rio de Janeiro, RJ, Brazil

The district of São Caetano (22° 03’08” S and 42° 41’17” W) arose after the colonization of the region of Vale do Rio Paquequer. The Downtown of São Caetano is 1 km from Portão de Pedra site, where the field collections were conducted with CDC-type traps, using light baits, in the cave, in the surroundings, and in the forest existing in the region. From now on, the cave will be called “cave of São Caetano”, for a better understanding of the texts. (Fig. 2)

**Fig 2.**
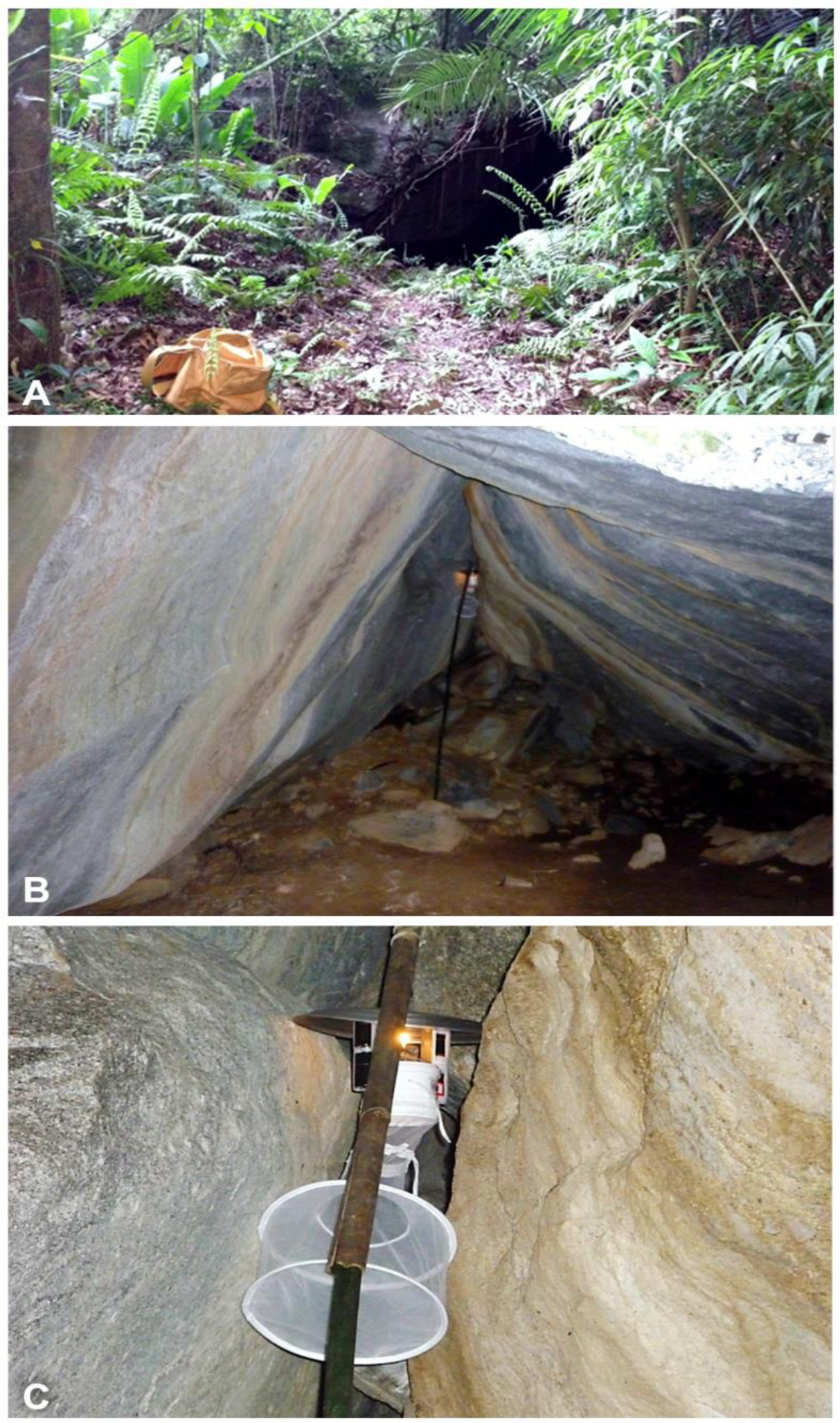
Entrance of the cave, near the base of São Caetano Rock, where the phlebotomine sand fly collections were made (A). CDC-type light trap between the rocks inside the cave (B). In detail, CDC-type light trap, between the rocks inside the cave (C). Photos of: Alves, JRC

The study area is located at the rock of São Caetano (22^a^ 03′02” S and 42° 42’07” W), which has 2 km in diameter and 500 meters in height, consisting of granite with vegetation coverage at its top, with a typical forest vegetation around it, located 3 km from the city headquarters.

### Collection of Specimens

In 2009 and 2010, captures were conducted with CDC light traps, HP model, twice a month, totaling 24 hours. Three luminous traps were used: one in the forest, one in the surroundings and one inside the cave, from 6 p. m to 6 a. m of the next morning. The climatic variables (temperature and relative humidity of the air), were provided by the National Institute of Meteorology according to its 6^th^ District of Meteorology – Observation and Applied Meteorology Section – SEOMA., Parque Nacional da Serra dos Órgãos, in the city of Teresópolis, Rio de Janeiro.

Considering the needs to verify the phlebotomine sand fly fauna profile, after the environmental changes resulting from the torrential rains of January 12, 2011, complementary captures were made in the same site, also with light traps and respecting the same systematization, from March 2015 to February 2016. The objective of these captures was to correlate the biodiversity and abundance of these insects and the possible vectors of leishmanias.

Captured phlebotomine sand flies were aspirated using manual suction catchers, were put at low temperature for 10 minutes and then transferred to a cylindrical tube with alcohol at 70%, properly labeled with the collection data and submitted to Diptera Laboratory, Phlebotomine Sector of Fundação Oswaldo Cruz, for sorting, arranging and identification.

For the classification and arrangement, the Young & Perkins (1984) technique was used, as modified by Aguiar (1993). The identification of the species of the *Lutzomyia* França, 1924 and *Brumptomyia* França & Parrot, genera was undertaken in accordance with Young & Duncan (1994) and Forattini (1973), respectively. (Fig 3.) The material is deposited in the Entomological Biodiversity Laboratory and in the Diptera Laboratory of the Oswaldo Cruz Institute, Oswaldo Cruz Foundation.

**Fig 3.**
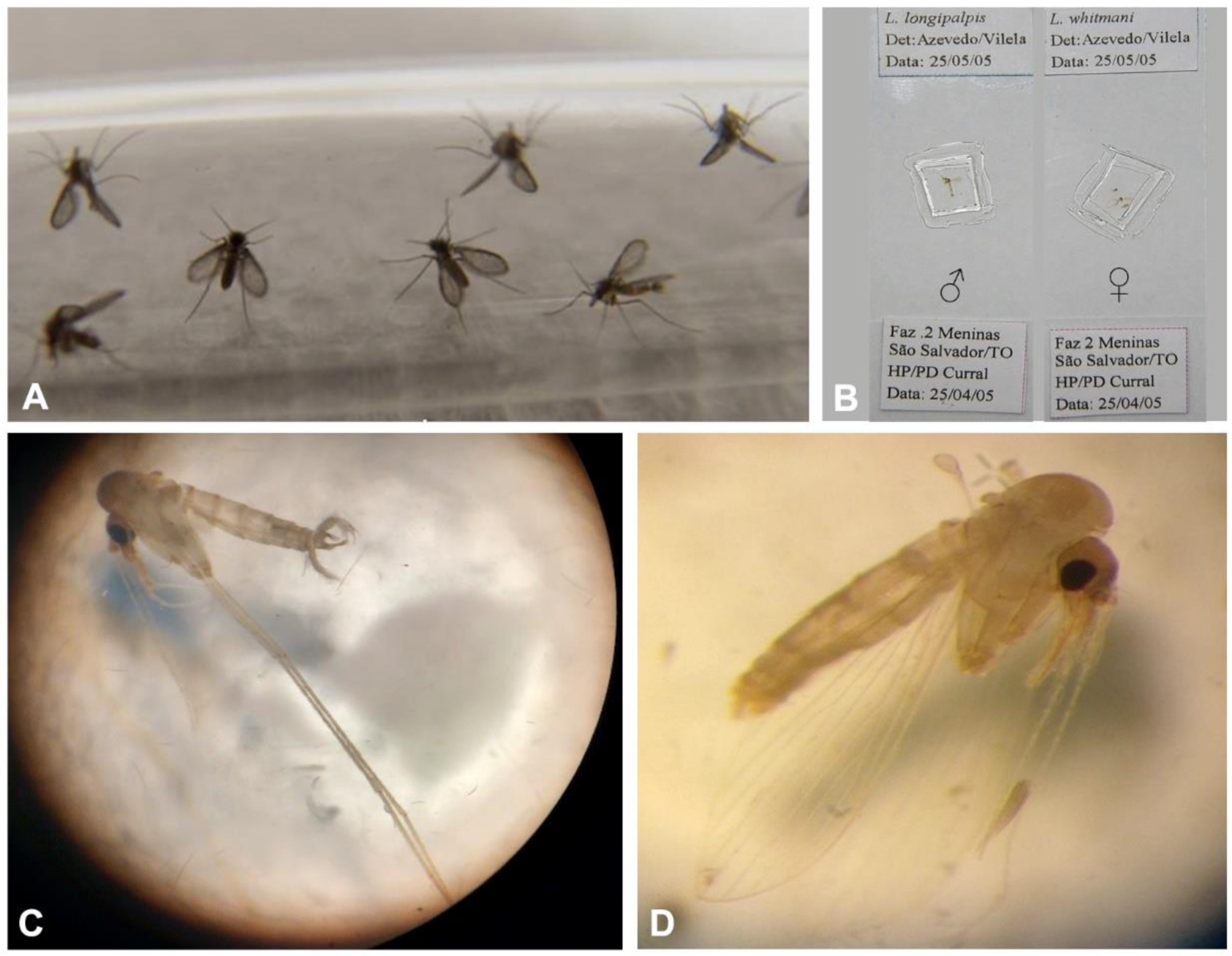
Male and female specimens of *Lutzomyia intermedia*, vector of *Leishmania braziliensis* in the State of Rio de Janeiro (A). Phlebotomine sand flies arranged in the Lamina and Glass slide in Berlese, Male (♂) and Female (♀) (B). Male (C) and female (D) phlebotomine sand flies, in the reagent, ready to be arranged in the lamina and glass slides in Berlese. Photos A and B of Vilela, ML; C and D of Alves, JRC.

### Statistical analysis

For the abundance and spatial distribution analysis of phlebotomines of a specific site, the Index of Species Abundance (ISA), the Standardized Index of Species Abundance (SISA) (Roberts and HSI 1979) was used.

The ISA was calculated in Microsoft Excel 2013 (Microsoft Corp., Redmond, WA, USA) and the values were converted between 0 and 1 (SISA), based on the following equations:

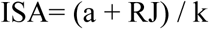

SISA= (c-ISA) / (c-1) where: K = capture number

a = value obtained by multiplying the number of the species absence (NAE) in k captures per c.

c = value of the highest position of the species in k captures plus 1.

RJ = sum of classifications in each species

For the results obtained in the research on the influence of temperature (C °) and relative humidity of air (RHA) on the phlebotomine fauna, in both periods, the Poisson Regression was used. Meanwhile, the ANOVA procedure was performed to compare the SISA results for *L. gasparviannai, L. edwardsi* and *L. tupynambai*.

### Results and discussion

From March 2015 to February 2016, 2.323 phlebotomines were captured at Sítio Portão da Pedra, belonging to nine species, with eight of *Lutzomyia* genus: *L. gasparviannai* (Martins, Godoy & Silva, 1962b), *L. edwardsi* (Mangabeira, 1946), *L. tupynambai* (Mangabeira, 1942b), *L. hirsuta* (Mangabeira, 1942b), *L. whitmani* (Antunes & Coutinho, 1939), *L. migonei* (França, 1920), *L. intermedia* (Lutz & Neiva, 1912) and *L*. sp; and one species of *Brumptomyia* genus: *Brumptomyia brumpti* (Larrouse, 1920) (Table 1).

**Table 1.**
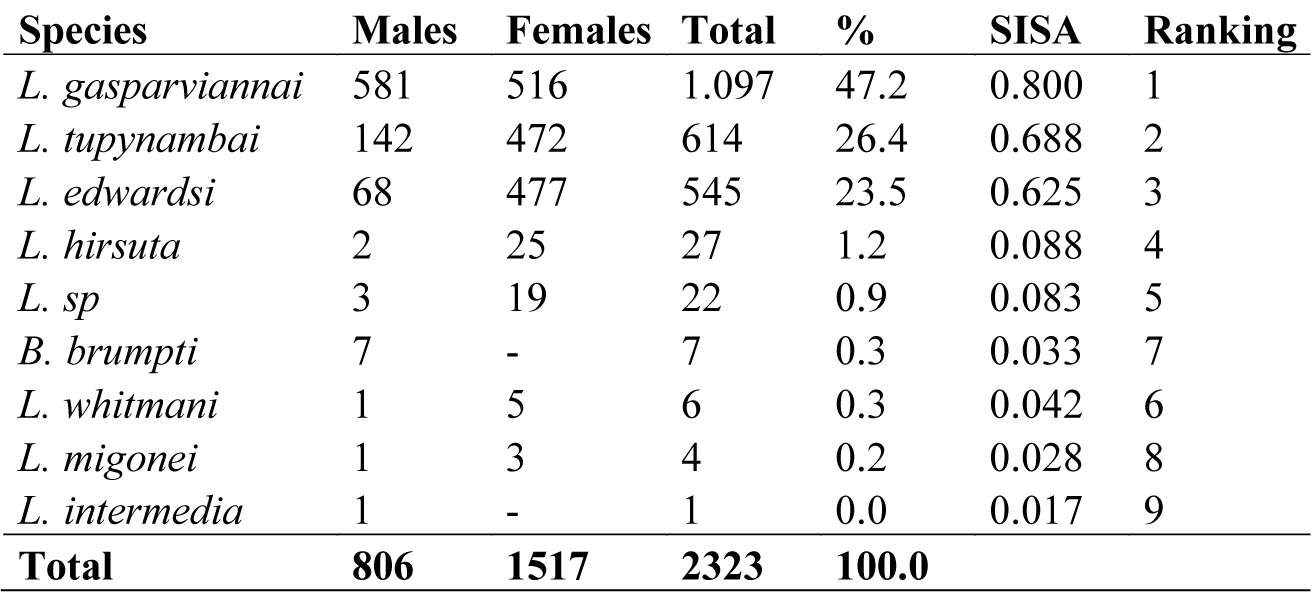
Frequency, Standardized Index of Species Abundance (SISA) and the total Rankin of phlebotomines captured with CDC model light trap placed in the cave, surroundings and Forest, from March 2015 to February 2016, at Sítio Portão de Pedra, in the city of Sumidouro, state of Rio de Janeiro, Brazil.

| Species | Males | Females | Total | % | SISA | Ranking |
| --- | --- | --- | --- | --- | --- | --- |
| *L. gasparviannai* | 581 | 516 | 1.097 | 47.2 | 0.800 | 1 |
| *L. tupynambai* | 142 | 472 | 614 | 26.4 | 0.688 | 2 |
| *L. edwardsi* | 68 | 477 | 545 | 23.5 | 0.625 | 3 |
| *L. hirsuta* | 2 | 25 | 27 | 1.2 | 0.088 | 4 |
| *L. sp* | 3 | 19 | 22 | 0.9 | 0.083 | 5 |
| *B. brumpti* | 7 | - | 7 | 0.3 | 0.033 | 7 |
| *L. whitmani* | 1 | 5 | 6 | 0.3 | 0.042 | 6 |
| *L. migonei* | 1 | 3 | 4 | 0.2 | 0.028 | 8 |
| *L. intermedia* | 1 | - | 1 | 0.0 | 0.017 | 9 |
| **Total** | **806** | **1517** | **2323** | **100.0** |  |  |

In 2009, the maximum temperature reached 30°C, with a minimum of 20°C, and in 2015 the maximum was 28.8°C and the minimum 17.7°C. The mean temperature was 23.7°C, for both periods and there was no variation.

Regarding the relative humidity of the air (RHA) there was a small difference, which can be explained by the heavy rains that fell on the region. The RHA average was 97% in 2015, while in 2009 it was 96.5%. (Fig. 4)

**Fig 4.**
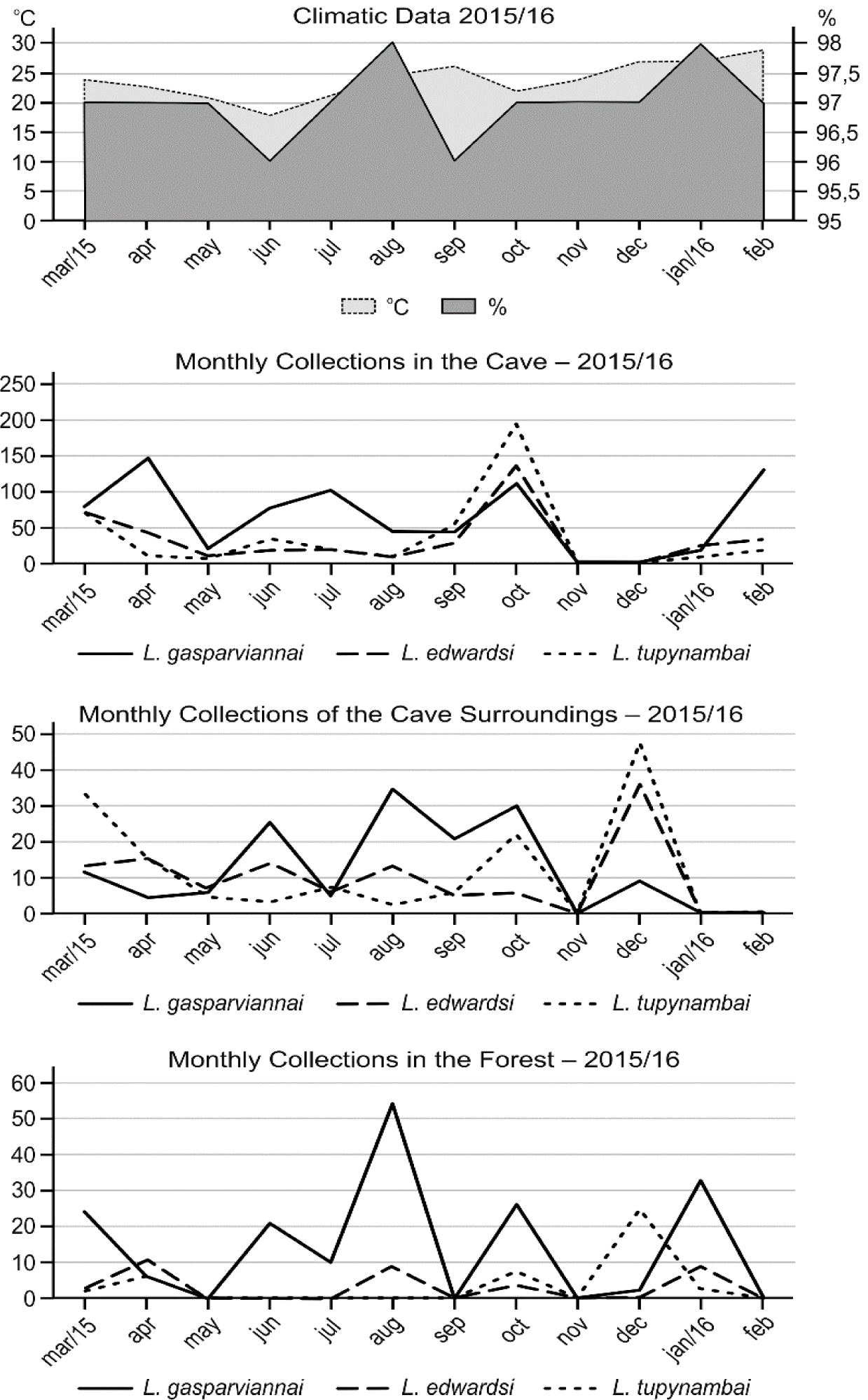
Total specimens of *L. gasparviannai, L. tupynambai* and *L. edwardsi* captured with CDC model light trap placed in the cave, forest and surroundings at Sitio Portão de Pedra, in the city of Sumidouro, state of Rio de Janeiro, Brazil, from March 2015 to February 2016, and its monthly frequency compared to the relative air humidity (%) and Temperature (°C)

The results of the Poisson regression showed that for *L. gasparviannai* and *L. edwardsi* when compared to temperatures (°C) of the collection periods of 2009/2010 and 2015/2016, there were no statistically significant differences (p-value = 0.328 and p-value=0.685, respectively), while for *L. tupynambai* significant statistical differences were found (p-value <0.01), with a negative sign, this means an opposite behavior between the variables. Considering the statistical method and the periods mentioned above, for *L. gasparviannai, L. edwardsi* and *L. tupynambai*, when comparing the relative humidity of the air (%) all these species showed statistically significant differences (p-value <0.01).

Considering the three collection sites, cave, surroundings and forest, we point out that:

*Lutzomyia gasparviannai* predominated over all species, types and collection sites, and 67.2% of females and 73.4% of males of this species were collected in the cave of São Caetano. *L. tupynambai* was the second most frequent species, with 67.5% of females and 79.6% of males captured in the cave, followed by *L. edwardsi* with 54.4% of males and 74.6% of females of this species in the cave. (Table 2).

**Table 2.** Total genders of phlebotomines captured with light traps (CDC) in the cave, surroundings and Forest, in the three collection sites, from March 2015 to February 2016, at Sítio Portão de Pedra, in the city of Sumidouro, state of Rio de Janeiro, Brazil.

| Species | Cave |  |  | Surrounding |  |  | Forest |  |  |
| --- | --- | --- | --- | --- | --- | --- | --- | --- | --- |
|  | Males | Females | Total | Males | Females | Total | Males | Females | Total |
| *L. gasparviannai* | 427 | 347 | 774 | 62 | 84 | 146 | 92 | 85 | 177 |
| *L. tupynambai* | 113 | 319 | 432 | 22 | 117 | 139 | 7 | 36 | 43 |
| *L. edwardsi* | 37 | 356 | 393 | 29 | 87 | 116 | 2 | 34 | 36 |
| *L. sp* | 3 | 18 | 21 | - | 1 | 1 | - | - | - |
| *L. hirsuta* | - | 17 | 17 | 1 | 7 | 8 | 1 | 1 | 2 |
| *B. brumpti* | 6 | 0 | 6 | - | - | - | 1 | - | 1 |
| *L. migonei* | - | 3 | 3 | 1 | - | 1 | - | - | - |
| *L. whitmani* | - | 3 | 3 | - | - | - | 1 | 2 | 3 |
| *L. intermedia* | - | - | - | 1 | - | 1 | - | - | - |
| **Total** | **586** | **1063** | **1649** | **116** | **296** | **412** | **104** | **158** | **262** |

We observed that *L. gasparviannai* was predominant, but the predominance of *L. edwardsi* females in the cave should be recorded, both in absolute numbers, 356 specimens, and frequency (74.6%), although it is the third in the general classification. In 2009 and 2010, 51.1% of females of this species were found in the cave, which increased considerably (Table 2). In 2016, only one male specimen was collected in the city of Cantagalo (Peres-Dias et al. 2016), and the literature points out that the climatic events that caused the tragedy of January 2011 (SDEISS 2011), did not reach the mentioned city. Considering these facts and the increase of this species in our study, we can suggest that there was a greater adaptation of this population compared to the previous and in the cities of Bom Jardim and Petrópolis, as recorded by Souza et al. (2002, 2005a, 2005) and De Souza et al. (2003).

Regarding *L. tupynambai* (79.6%), a difference in the frequency of the males of this species is pointed out compared to *L. gasparviannai* (73.4%), which does not confirms Rodrigues et al. (2013), who in their collection with CDC light trap, captured more females of *L. tupynambai* than males. At the same time, Aguiar & Vieira (2018) do not point out as one of the main habitats of these species, males and females, the rocky environments or crevices in the rock. For *L. tupynambai* they point out the armadillos’ burrow, other types of wild animals burrows, tree trunks, tubular roots, and marginal areas, while for *L. gasparviannai* only the forest with no defined location (Aguiar & Vieira 2018). This fact is relevant, because it is known that a considerable population of males on a given site suggests the existing of breeding sites of these phlebotomines (Alves 2007, Brazil and Brazil 2003). And such fact may have become necessary for the egglaying and, consequently, the perpetuation of the species due to sudden changes occurred over time.

Regarding the Standardized Index of Species Abundance (SISA), *L. gasparviannai* showed the highest abundance in both periods, slightly higher in the previous (0.708) and lower in this period (0.656); however, in absolute values, they were more significant in this study (774 phlebotomines) than in the previous (672 specimens), therefore showing a relevance of SISA indexes, because a more defined distribution suggests the need for control, prevention and research actions of this and other species for a longer time due to their homogeneity in space and time. Considering that, in addition to these, there is no record of *L. gasparviannai* in cave in the literature, with no parameter to discuss and analyze the real importance of these data, which will serve to compare future indexes. This is corroborated by Alves (2011), Barata et al. (2008, 2012), Campos et al. (2014), Carvalho et al. (2014) and Galati et al. (2010), (2003). (Table 3).

**Table 3.** Quantitative comparison of total, frequency, Standardized Index of Species Abundance (SISA) and the total Ranking of phlebotomines captured with CDC model light trap placed in the cave, surroundings and Forest, from June 2009 to May 2010, and from March 2015 to February 2016, at Sítio Portão de Pedra, in the city of Sumidouro, state of Rio de Janeiro, Brazil.

| Species | Total | % | SISA | Ranking | Total | % | SISA | Ranking |
| --- | --- | --- | --- | --- | --- | --- | --- | --- |
| Cave 2009 - 2010 |  |  |  |  | Cave 2015 - 2016 |  |  |  |
| *L. gasparviannai* | 672 | 66.3 | 0.708 | 1 | 774 | 46.9 | 0.656 | 1 |
| *L. tupynambai* | 159 | 15.7 | 0.476 | 2 | 432 | 26.2 | 0.508 | 2 |
| *L. edwardsi* | 151 | 14.9 | 0.316 | 3 | 393 | 23.8 | 0.490 | 3 |
| *L. hirsuta* | 9 | 0.9 | 0.122 | 4 | 17 | 1.0 | 0.021 | 6 |
| *L. whitmani* | 8 | 0.8 | 0.080 | 5 | 3 | 0.2 | 0.015 | 7 |
| *B. brumpti* | 4 | 0.4 | 0.049 | 6 | 6 | 0.4 | 0.025 | 5 |
| *L. migonei* | 3 | 0.3 | 0.042 | 7 | 3 | 0.2 | 0.010 | 8 |
| *L. sp.* | 1 | 0.1 | 0.017 | 10 | 21 | 1.3 | 0.072 | 4 |
| *L. microps* | 2 | 0.2 | 0.035 | 8 | - | - | - | - |
| *L. quinquefer* | 2 | 0.2 | 0.035 | 8 | - | - | - | - |
| *L. cortelezzii* | 2 | 0.2 | 0.028 | 9 | - | - | - | - |
| **Total** | **1013** | **100.0** |  |  | **1649** | **100.0** |  |  |
| Species | Surroundings 2009 - 2010 |  |  |  | Surroundings 2015 - 2016 |  |  |  |
| *L. gasparviannai* | 253 | 63.9 | 0.523 | 1 | 146 | 35.4 | 0.595 | 1 |
| *L. edwardsi* | 93 | 23.5 | 0.384 | 2 | 116 | 28.2 | 0.463 | 3 |
| *L. tupynambai* | 27 | 6.8 | 0.245 | 3 | 139 | 33.7 | 0.488 | 2 |
| *L. hirsuta* | 11 | 2.8 | 0.120 | 4 | 8 | 1.9 | 0.083 | 4 |
| *L. whitmani* | 4 | 1.0 | 0.056 | 5 | - | - | - | - |
| *L. davisi* | 2 | 0.5 | 0.046 | 6 | - | - | - | - |
| *L. migonei* | 2 | 0.5 | 0.028 | 7 | 1 | 0.2 | 0.017 | 5 |
| *B. sp.* | 2 | 0.5 | 0.023 | 8 | - | - | - | - |
| *L. sp.* | 1 | 0.3 | 0.019 | 9 | 1 | 0.2 | 0.012 | 6 |
| *B. guimaraesis* | 1 | 0.3 | 0.009 | 10 | - | - | - | - |
| *L. intermedia* | - | - | - | - | 1 | 0.2 | 0.017 | 5 |
| **Total** | **396** | **100.0** |  |  | **412** | **100.0** |  |  |
| Species | Forest 2009 - 2010 |  |  |  | Forest 2015 - 2016 |  |  |  |
| *L. gasparviannai* | 256 | 73.8 | 0.529 | 1 | 177 | 67.6 | 0.594 | 1 |
| *L. edwardsi* | 46 | 13.3 | 0.350 | 2 | 36 | 13.7 | 0.214 | 2 |
| *L. tupynambai* | 15 | 4.3 | 0.125 | 3 | 43 | 16.4 | 0.188 | 3 |
| *L. whitmani* | 12 | 3.5 | 0.108 | 4 | 3 | 1.1 | 0.083 | 4 |
| *L. hirsuta* | 11 | 3.2 | 0.088 | 6 | 2 | 0.8 | 0.083 | 4 |
| *L. davisi* | 3 | 0.9 | 0.063 | 5 | - | - | - | - |
| *L. sp.* | 3 | 0.9 | 0.050 | 8 | - | - | - | - |
| *B. brumpti* | 1 | 0.3 | 0.029 | 7 | 1 | 0.4 | 0.015 | 5 |
| **Total** | **347** | **100.0** |  |  | **262** | **100.0** |  |  |

A similar situation occurs with *L. tupynambai* that was higher in 2015/2016 with an index of 0.508, while in the previous it was 0.476. The absolute value between the periods is noteworthy, with a difference of 273 specimens between the species, although the index is not very distinct. These data do not corroborate the above mentioned authors.

*Lutzomyia edwardsi* showed a more homogeneous spatial distribution in 2015/2016, with 0.490 and a previous record of 0.316, which demonstrates a better and more efficient adaptation than the population of ten years ago (Table 3). Galati et al. collected *L. edwardsi* with an index of 0.353, and is the 6^th^ of the Ranking, with an absolute value of 48 phlebotomines considering the cave (Aguiar and Vieira 2018), outer wall of the cave (SDEISS 2011) and the forest (Azevedo and Rangel 1991, Galati et al. 2010). It was noticed that there was a more harmonic adaptation in the research conducted in this study, in Sumidouro. In both periods the species was the 3^rd^ of the Ranking, and the spatial distribution was more uniform, even after the environmental changes that occurred in the region. In 2015/2016, 242 more specimens were collected from a total of 544, almost half. (Table 3)

In both periods *L. edwardsi* was the third most frequent, and higher in this study with 23.8% of collected phlebotomines, while in the previous there were only 14.9% of phlebotomines, which confirms Campos et al. that in night collection in caves and surroundings in the city of Pains, recorded *L*. e*dwardsi* as the third most abundant (11%) and present in three cave environments (Campos et al. 2014). In absolute values, 151 and 393 specimens were collected in this study and in previous collection period in the cave of São Caetano, in Sumidouro, while Campos et al. (2014) collected 133 specimens in the cave, with 105 females and 28 males, and 30 specimens in the surroundings, with 25 females and 5 males. In that study, in 2009/2010, *L*. e*dwardsi* was the second in the Ranking of collections in the surroundings, with 93 phlebotomine, and six years later it was the third in the Ranking, but with 116 phlebotomines. In quantitative aspects, it is observed that *L*. e*dwardsi* was the most present in the cave of São Caetano and in its surroundings in Sumidouro, than in the research conducted by Galati et al. (2010) and Campos et al. (2014). The fact that in the city of Pains, the studied area suffers anthropic activity of mining, calcination and agriculture, may be contributing for the reduction of this fauna. Despite the landslides and erosions occurred in 2011 in São Caetano have been of great intensity, the fauna and flora have not suffered a massive and continuous anthropic activity since then. Alves (2007) reports the finding of *L. ayrosai, L. intermedia, L. lenti, L. migonei, L. quinquefer* and *L. whitmani* in a poorly diversified fauna in intradomiciliary, peridomiciliary and forest environments in a region with intense anthropic, pastoral and agricultural activity, where there was an autochthonous case of ATL in the city of Carmo. Ten years later, in the same place and with no intense anthropic activity, with the forest preserved, there is a rich fauna in species diversity (Carreira-Alves 2008). These facts demonstrate the ability of these phlebotomines to adapt to the environmental changes occurred over the years, both for natural and artificial causes, as recorded by Aguiar & Medeiros (2003), Aguiar & Vieira (2018), Carreira-Alves (2008) and Carvalho et al. (2014).

Poisson regression results showed that for *L. gasparviannai*, when compared with the collection periods of 2009/2010 and 2015/2016, no statistical differences were verified (p-valor=0.339); however, for *L. edwardsi* and *L. tupynambai* significant statistical differences were recorded (p-valor < 0.01).

The results of the Analysis of Variance (ANOVA) showed significant statistical evidence of differences in the SISA values between *L. gasparviannai, L. edwardsi* and *L. tupynambai* species (p-value = 0.004).

The monthly frequency of these species shows that *L. gasparviannai* had its highest peak in October (168), and it was decreasing until January, when it began to rise, showing a drop in March, subsequently recovering and stabilizing in the cold and dry period from June to August, and these months were responding for 33.9% of collected phlebotomines, while in the other months it was 27.1% more frequent. These data suggest a greater activity of this species in cold and dry places, either for food searches, mating or egglaying. In absolute values, in October there was the highest peak among males (108), and in April the females (92) were more significant. In 2009 and 2010, the most appropriate season were the warm and humid months, with 54.1% of collected phlebotomines, with a peak in March (240), and the female (191) was responsible for 75.6% of this total, with 20.4%. The lower temperature was 22°C and the highest humidity of the air was 98%, all in March. It is known that the higher the air humidity, more phlebotomines appear in nature (Aguiar and Medeiros 2003) (Table 4 and 5).

**Table 4.**
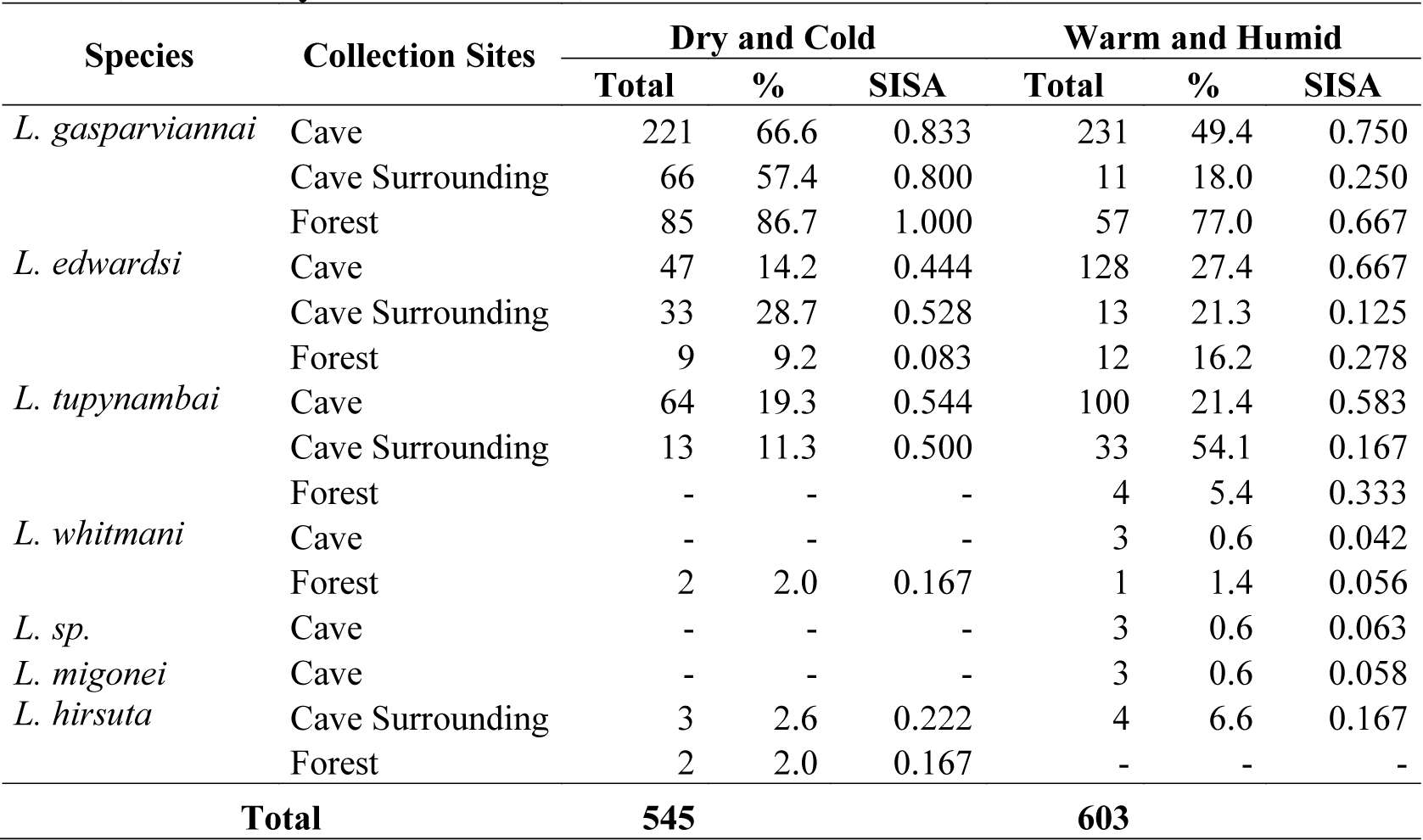
Frequency, Standardized Index of Species Abundance (SISA) and the total of phlebotomines captured with CDC model light trap placed in the cave, surroundings and Forest, in the cold and dry season, warm and humid season, from March 2015 to February 2016, at Sítio Portão de Pedra, in the city of Sumidouro, state of Rio de Janeiro, Brazil.

| Species | Collection Sites | Dry and Cold |  |  | Warm and Humid |  |  |
| --- | --- | --- | --- | --- | --- | --- | --- |
|  |  | Total | % | SISA | Total | % | SISA |
| *L. gasparviannai* | Cave | 221 | 66.6 | 0.833 | 231 | 49.4 | 0.750 |
|  | Cave Surrounding | 66 | 57.4 | 0.800 | 11 | 18.0 | 0.250 |
|  | Forest | 85 | 86.7 | 1.000 | 57 | 77.0 | 0.667 |
| *L. edwardsi* | Cave | 47 | 14.2 | 0.444 | 128 | 27.4 | 0.667 |
|  | Cave Surrounding | 33 | 28.7 | 0.528 | 13 | 21.3 | 0.125 |
|  | Forest | 9 | 9.2 | 0.083 | 12 | 16.2 | 0.278 |
| *L. tupynambai* | Cave | 64 | 19.3 | 0.544 | 100 | 21.4 | 0.583 |
|  | Cave Surrounding | 13 | 11.3 | 0.500 | 33 | 54.1 | 0.167 |
|  | Forest | - | - | - | 4 | 5.4 | 0.333 |
| *L. whitmani* | Cave | - | - | - | 3 | 0.6 | 0.042 |
|  | Forest | 2 | 2.0 | 0.167 | 1 | 1.4 | 0.056 |
| *L. sp.* | Cave | - | - | - | 3 | 0.6 | 0.063 |
| *L. migonei* | Cave | - | - | - | 3 | 0.6 | 0.058 |
| *L. hirsuta* | Cave Surrounding | 3 | 2.6 | 0.222 | 4 | 6.6 | 0.167 |
|  | Forest | 2 | 2.0 | 0.167 | - | - | - |
| **Total** |  | **545** |  |  | **603** |  |  |

**Table 5.** Monthly distributions of the total of *L. gasparviannai, L. tupynambai* and *L. edwardsi* captured with CDC model light tramp, compared with the temperature (T) and relative humidity of the air (RHA), from March 2015 to February 2016, at Sítio Portão de Pedra, in the city of Sumidouro, Rio de Janeiro, Brazil.

| Species | *L. gasparviannai* |  |  | *L. edwardsi* |  |  | *L. tupynambai* |  |  | T | RHA |
| --- | --- | --- | --- | --- | --- | --- | --- | --- | --- | --- | --- |
| Day/Mont<br>h | M | F | T | M | F | T | M | F | T | ° C | % |
| Mar-22 | 59 | 54 | 113 | 4 | 82 | 86 | 25 | 81 | 106 | 23.7 | 97 |
| Apr-22 | 65 | 92 | 157 | 8 | 60 | 68 | 3 | 27 | 30 | 22.6 | 97 |
| May-29 | 21 | 5 | 26 | - | 18 | 18 | 1 | 9 | 10 | 20.7 | 97 |
| Jun-20 | 44 | 80 | 124 | - | 32 | 32 | 17 | 20 | 37 | 17.7 | 96 |
| Jul-17 | 59 | 58 | 117 | 2 | 23 | 25 | 6 | 22 | 28 | 21.1 | 97 |
| Aug-15 | 88 | 43 | 131 | 10 | 23 | 33 | 3 | 9 | 12 | 24.6 | 98 |
| Sep-18 | 41 | 22 | 63 | 3 | 31 | 34 | 37 | 26 | 63 | 26.0 | 96 |
| Oct-28 | 108 | 60 | 168 | 23 | 122 | 145 | 30 | 195 | 225 | 21.8 | 97 |
| Nov-24 | - | - | - | - | - | - | - | - | - | 23.7 | 97 |
| Dec-14 | 6 | 6 | 12 | 10 | 27 | 37 | 12 | 60 | 72 | 26.8 | 97 |
| Jan-27 | 39 | 14 | 53 | 3 | 33 | 36 | 2 | 10 | 12 | 27.0 | 98 |
| Feb-9 | 50 | 82 | 132 | - | 31 | 31 | 6 | 13 | 19 | 28.8 | 97 |
| **Total** | **580** | **516** | **1,096** | **63** | **482** | **545** | **142** | **472** | **614** |  |  |

The monthly frequency shows that *L. edwardsi* had its highest peak in October (145), it was null in November, and started to rise in December (37), with a drop in January and February, recovering in March. In the warm and humid period it represented 28.7%, then it began to fall, with the lowest collection in May (18); from June to August fluctuated between 25 to 33 phlebotomines, and was higher in September (34), being responsible for 22.7% of collected phlebotomines. These data suggest a greater activity of this species in warmer period and in humid places, either for food searches, mating or egglaying. In absolute values, in October there was the highest peak among males (23), and females (122) were more significant. In 2009 and 2010, the most appropriate season were the warm and humid months, with 59.6% of collected phlebotomines, with a peak in March (46), and the females (42) were responsible for 91.3% of this total, and the male with 8.7%. The lower temperature was 22°C and the highest relative humidity of the air was 98%, all in March. (Fig. 4)

The monthly frequency of *L. tupynambai* shows that its peak was the highest of the three species and occurred in October (225); it was null in November, increased in December, and decreased from January to March, when there was a considerable peak (106). This period represented 22.3% of the total. In absolute values, in October there was the highest peak among males (195), while for females (37) it was in September. In 2009 and 2010, the most appropriate season were the warm and humid months, with 69.8% of collected phlebotomines, with a peak in March (106), and the females (85) were responsible for 80.1% of this total, and the male with 19.8%. The lower temperature was 22°C and the highest humidity of the air was 98%, all in March.

In the cave in 2009 and 2010, 11 species were collected, with 10 of *Lutzomyia* genus and one of *Brumptomyia* genus. In this study 8 species were collected, with 7 of *Lutzomyia* genus and one of *Brumptomyia* genus. There was a decrease in the diversity of species of the first genus, but *B. brumpti* occurred in both periods, confirming Aguiar & Vieira (2018), who pointed out that one of the main habitats of this species is the cave; however, the amount of specimens collected in that period and in this period are low, suggesting that there is a small population in the studied environment. This does not corroborate Alves (2008) and Carreira-Alves (2008), which in a study conducted in the city of Carmo, collected 30 specimens of *B. brumpti*, the second in frequency with 2.5% of the fauna species of the location.

According to Aguiar and Vilela (1987), the eating preferences of phlebotomines are a predominant factor, and directly influence their dispersion. The species of *Brumptomyia* genus, in its totality, suck blood from dasypodidae, and are always found in burrows of these animals and only accidentally out of them. Therefore, it can be said that where these animals do not occur, there is no representative of this genus (Aguiar and Medeiros 2003). This information was confirmed by Alves (2007), who conducted collections of phlebotomines in 1994 and 1995 in the city of Carmo, collected no *B. brumpti* and *B. cardosoi*, and has not recorded the occurrence of wild animals. But when he studied the same area in 2006 and 2007, it was found that in the collection site there were two burrows of dasypodidae with a radio of 3.65 m, which suggests that the appearance of these phlebotomines are due to the existence of armadillo burrows. These facts suggest the existence of dasypodidae in the cave of São Caetano and surroundings, therefore confirming the importance of knowing the food preference of phlebotomines while surveying its fauna, which dispersion can be observed with the collection of both in the studied area.

Alves (2008) and Carreira-Alves (2008) found that 33% of phlebotomines captured in the forest belonged to the *Brumptomyia* genus, confirming their silvatic behavior, as already observed by other authors (Aguiar and Medeiros 2003, Fraiha and Shaw 1970), which confirms the collection of *B. brumpti, B. guimaraesi* and *B*. sp in the forest and surroundings. We point out that the first species was collected in both periods, 2009/2010 and 2015/2016, in the cave of São Caetano and in the Forest, while the others only in the first period and in the surroundings (Table 6).

**Table 6.**
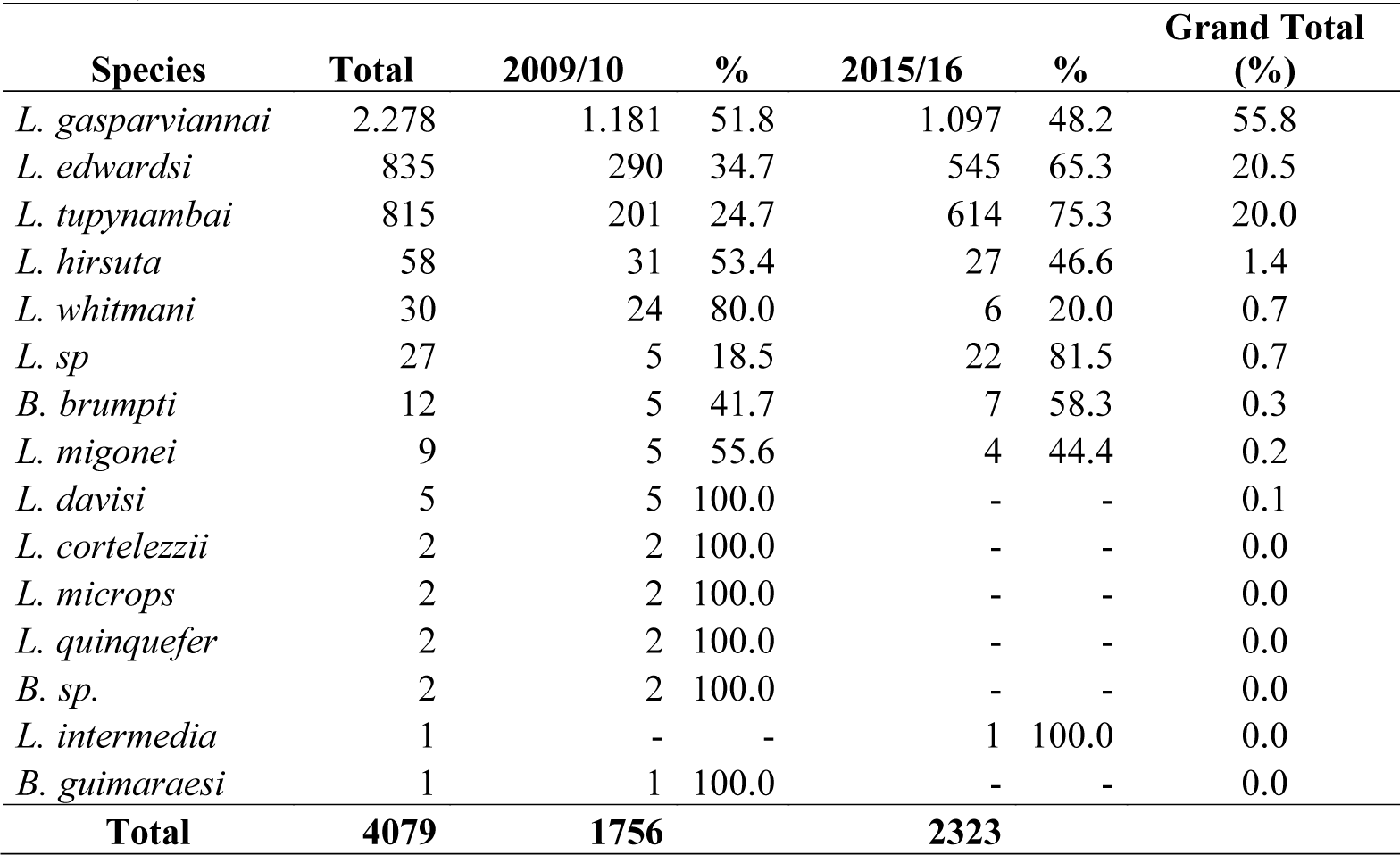
Quantitative comparison of total phlebotomines captured with CDC model light trap placed in the cave, surroundings and Forest, from June 2009 to May 2010, and from March 2015 to February 2016, at Sítio Portão de Pedra, in the city of Sumidouro, state of Rio de Janeiro, Brazil.

Despite the geographical distribution of the phlebotomine fauna of the above genus, *B. brumpti* was not recorded for the states of Rio de Janeiro and Espírito Santo by Aguiar & Medeiros (2003), and was later mentioned in the state of Rio de Janeiro for the cities of: Petrópolis, by Souza et al. (2005); Carmo, by Carreira-Alves (2008); Rio de Janeiro, by Gouveia et al. (2012), Carvalho et al.(2014); and Cantagalo, by Peres-Dias et al. (2016).

In both periods 23 specimens of *B. brumpti* were collected, with 2 males in the Forest, 2 females and 8 males in the cave of São Caetano. Campos et al. (2014), conducting the phlebotomine fauna study in caves of the city of Pains, state of Minas Gerais, has not recorded the finding of *B. brumpti* females, and at the same time recorded the collection of 5 males in the cave and 14 in the cave surroundings. This confirms the finding of Galati et al. (2003) when researched the cave fauna of Serra da Bodoquena in Mato Grosso do Sul, and corroborating with Aguiar & Vieira (2018), the authors do not point out this species to the state of Goiás, in the Midwest region of Brazil.

According to Carvalho et al. (2014) this is the second record of *B. brumpti, B. cardosoi* and *B. guimaraesi* for the mountain region of the state of Rio de Janeiro, and the first was made by Souza (2002, 2005). We point out that *B. avellari* and *B. nitzulescui* were mentioned for the first time in the city of Carmo and in the mountain region of this state by Carreira-Alves (2008), showing that from the eight species of *Brumptomyia* genus, five have already been recorded here. In this study, this diversity was restricted to 25% of the fauna known in this state.

*Lutzomyia* (*Psychodopygus*) *hirsuta* (Mangabeira, 1942) was collected in three collection sites, and in quantitative values it was greater and significant in the cave of São Caetano (17) and less in the Forest (2), respectively, in the second study period. In 2009/2010, it was the 4^th^ in the Ranking in the cave of São Caetano and its surroundings, and the 5^th^ in the Forest. Six years later it was the 4^th^ in the Ranking in the surrounding, in the Forest and the 6^th^ in the cave of São Caetano. This is the second time that this species is recorded in the mountain region of the state of Rio de Janeiro, and was mentioned 57 years ago by Martins (1962a, 1978) in the city of Petrópolis. The finding in two different ecosystems, Forest and cave, in both periods, quite shows its adaptation to the studied region. This fact is relevant considering that in 1985, Rangel et al. isolated *Leishmania* braziliensis of *L. hirsuta* in a forest close to the city of Além Paraíba, state of Minas Gerais, location type of *Leishmania* (*Viannia*) *braziliensis*. Recently, Gil et al. (2003) found *L. davisi* and *L. hirsuta* infected with *Leishmania* (*Viannia*) *naiffi* in the state of Rondônia. In 2019, Miranda recorded an autochthonous case of leishmaniasis in a rural area of the city of Duas Barras-RJ, with a genetic variant of *Leishmania* (*Viannia*) *naiffi*.

We point out that *L. davisi* and *L. hirsuta* are mentioned for the second time in the mountain region of the state of Rio de Janeiro, and the first species was collected in the city of Carmo by Carreira-Alves (2008). Considering the presence of phlebotomines in the cold and dry months (June, July and August) and warm and humid months (January, February and March), and no presence of *L. hirsuta* was recorded in the cave of São Caetano in the mentioned months in 2015/2016. In the cold and dry period the following temperatures were recorded, by order of the month: June with 17.7°C, July with 21.1°C and August with 24.6°C; while in the warm and humid months, they were: January with 27°C, February with 28.8°C and March with 23.7°C. Regarding the relative humidity of the air in the cold and dry season, 96%, 97% and 98% were obtained, while in the warm it was 98%, 97% and 97%. According to Aguiar & Vieira (2018), in the dry season the shelters temperature is greater than in the external environment and increases in the rainy season. The relative humidity of the air in the shelter, both in the cold and dry and in the warm and humid periods is always greater than the external environment. When the temperature change occurs gradually, and the air humidity is abrupt, the phlebotomines appear. The air humidity is the most important factor. Therefore, it can be noticed that the temperature change in these periods was not enough to significantly change the internal and external temperature of shelters, and also, the air humidity had no significant change, with indexes very close, which justify the absence of phlebotomines of this species.

In 2009 and 2010, *L. whitmani* was collected in the three collection sites, with 12 specimens in the Forest, 8 in the cave of São Caetano and 4 in the surrounding, representing 1.4% of the collected species, being the 5^th^ in the total Ranking. Regarding the gender, 22 females were confirmed and only 2 males. In 2015 and 2016, three specimens were collected in the cave of São Caetano and three in the Forest, with a very low frequency (0.3%), and considering its spatial distribution it was the 6^th^ in the total Ranking. Regarding the gender, the female was the most captured (5) again and only one male was found (Table 3). Rangel & Lainson (1990) considered *L. whitmani* a new wild species. However, it could be found inside homes that were in the middle of the forest or in neighboring areas. This was shared by Oliveira (2003), and in addition, they believed that *L. whitmani* was in the adaptation process to the anthropic environment in other areas of Brazil (Azevedo and Rangel 1991, Azevedo et al. 1996). In São Caetano it is common to find houses near the forest, and also one person had ATL, which lived in the margins of this site. Aguiar & Vieira (2018) pointed out in their study that *L. whitmani* can be found in the internal and external walls of human domicile, and in facilities of domestic animals (chicken coop, pigsty, corral etc.). They are also found in trunks, hollows and treetops, and in tubular roots. Although it is found in Sumidouro with a low frequency, which corroborates Alves (2007), in his study in Carmo, and such result was confirmed by Alves (2008). These data are relevant, because Ready et al. (2018) confirmed *L. whitmani* and *L. intermedia* as a vector of *Leishmania* (*Viannia*) *braziliensis* for the Northeast and Southeast, and *L. whitmani* in the Midwest of Brazil. They also confirmed that they have found *L. whitmani* infected with *Leishmania* (*Viannia*) *guyanemsis* and *Leishmania* (*Viannia*) *shawi* in the Amazon.

According to De Souza et al. (1995), Alves (2007) and Carreira-Alves (2008) the species *L. intermedia, L. whitmani*^*1*^, *L. migonei*^2^, *L. fischeri*^2^, *L. quinquefer*^2^ and *L. monticola*^2^ were recorded in the cities of Carmo and São José do Vale do Rio Preto, and also the five first species were recorded in Sumidouro. The species below were only collected in Carmo: *L. ayrozai*^2^, *L. aragaoi*^1^, *L. carrerai*^1^, *L. cortelezzii*^1^, *L. davisi*^1^, *L. lanei*^2^, *L. lenti*^1^, *L. lutziana*^2^, *L. sordellii*^1^, *B. avellari*^1^, *B. brumpti*^*2*^, *B. cardosoi*^2^, *B. guimaraesi*^2^ and *B. nitzulescui*^1^ (Carreira-Alves 2008). Meanwhile, *L. alencari*^*1*^, *L. edwardsi*^*2*^, *L. firmatoi*^*1*^ and *L. pelloni*^*1*^ were found in the city of São José do Vale do Rio Preto (De Souza et al.1995).

Except the cities of Carmo, São José do Vale do Rio Preto and Sapucaia, the mentioned species were not found in three cities, which are: Duas Barras, Nova Friburgo, and Teresópolis, which border Sumidouro. Martins et al. (1978) and Carvalho et al. (2014), recorded the finding of *L. schreberi* in the city of Teresópolis.

The species marked with (^1^) and (^2^) were recorded in the first and second time, in different cities of the mountain region of the state of Rio de Janeiro, considering De Souza et al. (1995), Carreira-Alves (2008) and Carvalho et al. (2014).

In the two periods of this study, *L. migonei* was present in the cave of São Caetano and in the surrounding of this cave, with only 3 specimens in each of the researched periods in the cave of São Caetano, with a very low frequency, and we can notice that its spatial distribution was higher in 2009/2010, with a SISA of 0.042, while in the second it was only 0.010. Regarding the genders, the presence of males and females was recorded in the cave of São Caetano in the first study, and in 2015/2016 only three females were recorded. Aguiar & Vieira (2018) have not pointed out the caves as one of the main habitats of this species. However, in the warm and humid periods it was present in the cave of São Caetano in 2015/2016. In the previous study, it was present both in cold and dry and in the warm and humid periods in the cave of São Caetano and its surroundings. These data suggest that the population existing, both in the cave and its surroundings, are low and showed that they have suffered a significant impact due to the heavy rains that affected in the region.

Considering that the air humidity is a major factor for the choice of habitats, feeding, egglaying and the appearance of phlebotomines, when its change is abrupt, it affects the biological cycle of these dipterans. They did not occur in these collections and the air humidity and temperature are close. The peaks were more intense in this study than in the previous, both in frequency and in absolute value. This shows that these species are well adapted to the two collection sites (cave of São Caetano and Forest).

In 2009 and 2010, *L. cortelezzii, L. microps* and *L. quinquefer* were only collected in the cave, *L. davisi, L. whitmani* and *B. guimaraesi* in the surroundings, and *L. davisi* in the Forest, and in 2015 and 2016, *L. intermedia* was collected in the surroundings. The *L. davisi, L. cortelezzii, L. microps, L. quinquefer, L. intermedia* and *B. guimaraesi* species were collected only once in the two researched periods, with low frequency and low absolute values. Therefore, it is not possible to make a correlation between them, because there is no parameter for that. It is believed that the abrupt changes occurred in 2011 did not allow the adaptation and survival of these species until 2015 and 2016 (Table 6).

The literature points out that in the collections with CDC-type light trap in the forest, the frequency of *L. intermedia* is low, which corroborates with the finding of this study (Alves 2007).

## Conclusions

Only future studies in the region could define the sand fly fauna of the above-mentioned species.

It is also verified that the changes have influenced the dispersion and distribution of the phlebotomine sand fly fauna existing in both ecosystems (cave of São Caetano and Forest). And also, they created ecological conditions that allowed the increase of populations more resistant to sharp weather events.

Due to this, it was observed that in 2015 and 2016 the phlebotomine sand fly fauna was more quantitative, but less diverse and with the presence of *L. intermedia, L. whitmani* and *L. migonei*, vectors *Leishmania*, but in small populations. The record of 31 cases of ATL, considering the city of Sumidouro and those from surroundings, from 2007 to 2017, the finding of these three species becomes a relevant data, which should deserve from the health authorities of the mountain region a greater attention regarding the surveillance and control of leishmaniasis vectors.

## ACKNOWLEDGEMENTS

To Dr. Felipe Ferraz Figueiredo Moreira, head of the Laboratory of Entomological Biodiversity of Fundação Oswaldo Cruz, for the support that was of vital importance for conducting this study.

To Dr. Márcio Félix of the Laboratory of Entomological Biodiversity of Fundação Oswaldo Cruz, for the help and support.

To Dr. Francisco Gerson de Araújo, Coordinator of the Postgraduate studies in Animal Biology of the Universidade Federal Rural of Rio de Janeiro, for the suggestions and advices.

The Heloisa Maria Nogueira Diniz the production and processing of images. This study was developed with the support of the Coordination for the Improvement of Higher Education Personnel – Brazil (CAPES) – Funding Code 001.

## CONFLICT OF INTEREST

The authors declare there is not conflict of interest.

## Notes

### Competing Interest Statement

The authors have declared no competing interest.

## REFERENCES CITED

Aguiar GM & Vieira VR. 2018. Regional distribution and habitats of Brazilian Phlebotomine Species. In: Rangel EF, Shaw JJ. (eds), Brazilian Sandflies. Rio de Janeiro, Springer, 489 p.

Aguiar GM, Vilela ML, Schuback, Soucasaux P, Azevedo ACR. 1985. Aspecto da ecologia dos flebótomos do Parque Nacional da Serra dos Órgãos, Rio de Janeiro III. Frequência horária (Diptera, Psychodidae, Phlebotominae). Mem Inst Oswaldo Cruz 80: 339–348.

Aguiar G M and Vilela ML. 1987. Aspects of the ecology of sandflies at the Serra dos Órgãos National Park, State of Rio de Janeiro. VI. Shelters and breeding places (Diptera, Psychodidae, Phlebotominae). Mem Inst Oswaldo Cruz 82:585–586.

Aguiar GM. 1993. Estudo sobre a ecologia dos flebotomíneos da Serra do Mar, município de Itaguaí, estado do Rio de Janeiro, Brasil, área de transmissão de leishmaniose tegumentar (Diptera: Psychodidae, Phlebotominae). Ph. D. Tese, Universidade Federal do Paraná, Curitiba.

Aguiar GM, Medeiros WM. 2003. Distribuição regional e hábitats das espécies de flebotomíneos do Brasil. In: Rangel EF, Lainson R. (eds.) Flebotomíneos do Brasil. Rio de Janeiro. Fundação Oswaldo Cruz, 368 p.

Alves JRC. 2007. Espécies de *Lutzomyia* França (Diptera: Psychodidae, Phlebotominae) em Área de Leishmaniose Tegumentar no Município de Carmo, RJ. Neotrop Entomol 36: 593–596.

Alves JRC. 2008. Espécies de Phlebotominae (Diptera: Psychodidae) da fazenda São José, Município de Carmo, estado do Rio de Janeiro, Brasil. Dissertação de Mestrado. Universidade Federal Rural do Rio de Janeiro, Seropédica.

Alves VR, Freitas RA, Santos FL, Barrett TV. 2011. Diversity of sandflies (Psychodidae: Phlebotominae) captured in sandstone caves from Central Amazonia, Brazil. Mem Inst Oswaldo Cruz 106: 353–359.

Azevedo ACR and Rangel EF. 1991. A study of sandfly species (Diptera: Psychodidae: Phlebotominae) in a focus cutaneous leishmaniasis in the Municipality of Baturité, Ceará, Brazil. Mem Inst Oswaldo Cruz.; 86: 405–410.

Azevedo ACR, Vilela ML, Souza NA, Andrade-Coelho CA, Barbosa AF, Firmino ALS, et al. 1996. The sandfly fauna (Diptera, Psychodidae: Phlebotominae) of a focus of cutaneous leishmaniasis in Ilhéus, State of Bahia, Brazil. Mem Inst Oswaldo Cruz 91: 75–79.

Barata RA, Antonini Y, Macedo CG, Costa DC, Dias ES. 2008, Flebotomíneos do Parque Nacional Cavernas do Peruaçu, Minas Gerais, Brasil. Neotrop Entomol 37: 226–228.

Barata RA, Serra e Meira PCL, Carvalho GML. 2012. *Lutzomyia diamantinensis* sp. nov., a new phlebotomine species (Diptera, Psychodidae) from a quartzite cave in Diamantina, Minas Gerais State, Brazil. Mem Inst Oswaldo Cruz 107: 1006–1010.

Brazil RP and Brazil BG. 2003. Biologia de flebotomíneos do Brasil, In: Rangel EF, Lainson R, (eds.) Flebotomíneos do Brasil. Rio de Janeiro, Fundação Oswaldo Cruz. 367 p.

Campos A M, Dos Santos CLC, Stumpp R, Da Silva LHD, Maia RA, Paglia AP, et al. 2017. Photoperiod Differences in Sand Fly (Diptera: Psychodidae) Species Richness and Abundance in Caves in Minas Gerais State, Brazil. J Med Entom 54: 100–105.

Carreira-Alves JR. 2008. Espécies de Phlebotominae (Diptera: Psychodidae) da fazenda São José, Município de Carmo, estado do Rio de Janeiro, Brasil. Rev Patol Trop 37: 371–372.

Carvalho BM, Dias CM, Rangel EF. 2014. Phlebotomine sand flies (Diptera, Psychodidae) from Rio de Janeiro State, Brazil: species distribution and potential vectors of *leishmaniasis*. Rev Bras Ent 58: 58 – 87.

Carvalho GMDL, Brazil RP, Ramos MCDNF, Serra e Meira PCL, Zenóbio APLDA, Botelho HA, et al. 2013. Ecological aspects of phlebotomine sandflies (Diptera: Psychodidae) from a cave of the speleological province of Bambuí, Brazil. PloS ONE 8: 10. 9 pages.

De Souza MB, Marzochi MCA, Carvalho RW, Conceição NF, Pontes CS. 1995. Flebótomos em áreas de ocorrência de leishmaniose tegumentar no Município de São José do Vale do Rio Preto, Rio de Janeiro, Brasil. Parasitol al Dia (Flap) 97 – 103.

De Souza MB, Cardoso PG, Sanavria A, Marzochi MCA, Carvalho RW, Ribeiro PC et al. 2003. Fauna Flebotomínica do município de Bom Jardim, região serrana do estado do Rio de Janeiro, Brasil. Revista Brasileira de Parasitologia Veterinária 12: 150 – 153.

Foratini OP. 1960. Novas observações sobre a biologia de flebótomos em condições naturais (Diptera: Psychodidae). Arq Hig Saúde Publica 25: 209–215.

Fraiha H and Shaw JJ, R. 1970. Phlebotominae brasileiros 1: Descrição de uma espécie de *Brumptomyia* e chave para identificação dos machos das espécies do gênero (Diptera: Psychodidae). Rev Bras Biol 30: 465–470.

Galati EAB. 2003. Morfologia, terminologia de adultos e identificação dos táxons da América, In: Rangel EF, Lainson R, (eds.) Flebotomíneos do Brasil. Rio de Janeiro, Fundação Oswaldo Cruz. 367 p.

Galati EAB, Marassá AM, Gonçalves-Andrade RM, Consales CA, Bueno EFM. 2010. Phlebotomines (Diptera, Psychodidae) in the Ribeira Valley Spelological Province1. Parque Estadual Intervales, State of São Paulo, Brazil. Rev Bras Ent 54: 311–321.

Gil LHS, Basano SA, Souza AA, Silva MGS. Barata I, Ishikawa EA, et al. 2003. Recent observations on the sand fly (Diptera: Psychodidae) Fauna of the state of Rondônia, Westem Amazonia, Brazil: The importance of *Psychodopygus davisi* as a vector of zoonotic cutaneous leishmaniasis. Mem. Inst Oswaldo Cruz 98: 751–755.

Gouveia C, Oliveira RM, Zwetsch A, Motta-Silva D, Carvalho BM, Santana AF, et al. 2012. Integrated tools for American cutaneous leishmaniasis surveillance and control: Intervention in an endemic area in Rio de Janeiro, RJ, Brazil. Inte Pers Infectious Diseases, 9 pages

Lima LC, Marzochi MCA, Sabroza PC, Souza MA. 1988. Observação sobre a leishmaniose tegumentar, cinco anos após profilaxia. Rev Saúde Pública 22: 73–77.

Martins AV, Godoy Jr TL, Silva JE. 1962a. Nota sobre os flebótomos de Petrópolis, estado do Rio de Janeiro, com a descrição de uma nova espécie (Diptera, Psychodidae). Rev Bras Biologia 22: 55 – 60.

Martins AV, Williams P, Falcão AL. 1978. American Sandflies (Diptera, Psychodidae, Phlebotominae). Academia Brasileira de Ciências, Rio de Janeiro, RJ. 195 p.

Menezes CRV, Azevedo ACR, Costa SM, Costa WA, Rangel EF. 2002. Ecology of American *Leishnaniasis* in the State of Rio de Janeiro, Brazil. J Vec Ecology 27: 207–214.

Miranda LFC. 2019. Caracterização taxonômica e distribuição espacial das espécies causadoras de Leishmaniose tegumentar na população humana e canina atendida no Instituto Nacional de Infectologia Evandro Chagas, de 2000 a 2015 [tese]. Fundação Oswaldo Cruz, Rio de Janeiro.

Oliveira AG, Andrade Filho JD, Falcão AL, Brazil RP. 2003. Estudo de flebotomíneos (Diptera, Psychodidae, Phlebotominae) na zona urbana da Cidade de Campo Grande, Mato Grosso do Sul, Brasil, 1999-2000. Cad Saúde Pública19: p. 933 – 944.

Peres-Dias QN, Oliveira CD, Souza MB, Meira AM, Villanova CB. 2016. S and fly species composition (Diptera: Psychodidae: Phlebotominae) in the municipality of Cantagalo, an area with sporadic cases of human cutaneous Leishmaniasis in Rio de Janeiro state, Brazil. Rev Inst Med Trop São Paulo [Internet]. [cited 2020 Jan 15]:58:50.

Prefeitura Municipal de Sumidouro (PMS): informações sobre o meio ambiente, saúde, educação, segurança. [Disponível em: http://sumidouro.rj.Gov.br/]. [Acessado em 17 abril 2019]

Rangel EF, Ryan L, Lainson R, Shaw JJ. 1985. Observations on the sandfly (Diptera: Psychodidae) fauna of Além Paraíba, state of Minas Gerais, Brazil, and the isolation of a parasite of the *Leishmania braziliensis* complex from *Psychodopygus hirsuta hirsuta*. Mem. Inst. Oswaldo Cruz 80: 373–374.

Rangel EF, Azevedo ACR, Andrade CA, Souza NA, Wermelinger ED. 1990. Studies on sandfly fauna (Diptera: Psychodidae) in focus of cutaneous leishmaniasis in Mesquita, Rio de Janeiro State, Brazil. Mem Inst Oswaldo Cruz 85: 39–45.

Rangel EF and Lainson R. 2003. Ecologia das Leishmanioses. In: Rangel, E.F, Lainson, R. (eds.) Flebotomíneos do Brasil, Editora FIOCRUZ, 489 p.

Ready PD, Vigoder FM, Rangel EF. 2018. Molecular and Biochemical Markers for Investigating the Vectorial Roles of Brazilian Sandflies. In: Rangel E.F, Shaw J.J. Editors, Brazilian Sandflies. Rio de Janeiro, Springer. 489 p.

Roberts DR and His BP. 1979. An index of species abundance for use with mosquito surveillance data. Env Entomology 8: 1007–1013.

Rodrigues AAF, Barbosa VA, Andrade-Filho JD and Brazil RP. 2013. The sandfly fauna (Diptera: Psychodidae: Phlebotominae) of the Parque Estadual da Serra da Tiririca, Rio de Janeiro, Brazil. Mem Inst Oswaldo Cruz 108: 943–946.

Secretaria de Desenvolvimento Econômico, Energia, Indústria e Serviço-RJ (SDEEIS). 2011. Megadesastre da Serra, Janeiro de 2011. Departamento de Recursos Minerais (DRM)/RJ.

Souza NA, Andrade-Coelho CA, Peixoto AA, Rangel EF. 2005. Nocturnal activity rhythms of *Lutzomyia intermedia* and *Lutzomyia whitmani* (Diptera: Psychodidae) in a transmission area of American cutaneous leishmaniasis in Rio de Janeiro State, Brazil. J Med Ent 42: 986–992.

Souza NA, Andrade-Coelho CA, Vilela ML, Peixoto A, Rangel EF. 2002. Seasonality of *Lutzomyia intermedia* and *Lutzomyia whitmani* (Diptera: Psychodidae: Phlebotominae), occurring sympatrically in area of Cutaneous Leishmaniasis in the State of Rio de Janeiro, Brazil. Mem Inst Oswaldo Cruz 97: 759–765

Souza MB, Cardoso PG, Sanavria A, Marzochi MCA, Carvalho RW, Ribeiro P.C, et al. 2003a. Fauna flebotomínica do município de Bom Jardim, região serrana do estado do Rio de Janeiro, Brasil. Rev Bras Paras Vet 12: 150 –153.

Souza NA, Andrade-Coelho CA, Silva VC, Peixoto AA, Rangel EF. 2005. Moonlight and blood-feeding behavior of *Lutzomyia intermedia* and *Lutzomyia whitmani* (Diptera: Psychodidae: Phlebotominae), vectors of American cutaneous leishmaniasis in Brazil. Mem Inst Oswaldo Cruz 100: 39–42

Teodoro U, et al. 1999. Impacto de alterações ambientais na ecologia de flebotomíneos no sul do Brasil. Cad Saúde Pública.; 15: 901–906.

Young DG and Duncan MA. 1994. Guide to the identification and geographic distribution of *Lutzomyia* sand flies in Mexico, the west Indies, Central and Douth American (Diptera: Psychodidae). 1o ed. Memories of the American Entomological Institute.

Young DC, and Perkins PV. 1984. Phlebotominae Sand Flies of North America (Diptera: Psychodidae). Mosq News 44: 263–304

